# Resting-State Network Analysis of Suicide Attempt History in the UK Biobank

**DOI:** 10.1101/2022.01.04.474947

**Authors:** Matthew F. Thompson, Marjan Ghahramanlou-Holloway, Mikela A. Murphy, Kanchana U. Perera, Chelsie Benca-Bachman, Rohan H. C. Palmer, Joshua C. Gray

## Abstract

**Background:** Prior research has identified altered brain structure and function in individuals at risk for self-directed violence thoughts and behaviors. However, these studies have largely utilized healthy controls and findings have been inconsistent. Thus, this study examined differences in resting-state functional network connectivity among individuals with lifetime suicide attempt(s) versus lifetime self-directed violence thoughts alone.

**Methods:** Using data from the UK Biobank, this study utilized a series of linear regressions to compare individuals with lifetime suicide attempt(s) (*n* = 566) versus lifetime self-directed violence thoughts alone (*n* = 3,447) on within- and between-network resting-state functional connectivity subnetworks.

**Results:** There were no significant between-group differences for between-network, within-network, or whole-brain functional connectivity after adjusting for age, sex, ethnicity, and body mass index and performing statistical corrections for multiple comparisons.

**Conclusions:** Resting-state network measures may not differentiate between individuals with lifetime suicide attempt(s) and lifetime self-directed violence thoughts alone. Null findings diverge from results reported in smaller neuroimaging studies of suicide risk, but are consistent with null findings in other large-scale studies and meta-analyses. Strengths of the study include its large sample size and stringent control group. Future research on a wider array of imaging, genetic, and psychosocial risk factors can clarify relative contributions of individual and combined variables to suicide risk and inform scientific understanding of ideation-to-action framework.

## Introduction

For each attempted suicide, it is estimated that 7-8 individuals have thoughts of suicide without attempting^1-3^. Prevailing theories on the ideation-to-action framework of suicide conceptualize distinct etiologic factors leading to the development of suicidal thoughts versus suicidal behaviors^4^. While there is increasing empirical support for distinct etiologic factors^4-6^, biological correlates of suicidal thoughts alone versus behaviors remain an understudied area in suicide prevention^7-12^. Exploring biological correlates of suicide risk, especially through means like neuroimaging, provides insight into potential differences between those who only think about suicide versus those who subsequently progress from suicidal ideation to suicidal action(s)^12^.

Imaging studies have found differences among individuals with versus without lifetime suicide-related thoughts and behaviors (STBs), though specific findings vary widely by control group, sample size, and imaging methods^13^. Recent meta-analyses have shown differences in brain activation when comparing suicidal individuals to healthy controls^9, 14^, but few consistent differences have emerged among studies using psychiatric or suicide-related controls. Based on a meta-analysis relying on data from 533 individuals from both task and resting-state modalities, Huang and colleagues (2020)^9^ found hyperactivation in the temporoparietal junction among those with lifetime STBs when compared to psychiatric controls (i.e., those without STBs but with psychiatric diagnosis or clinical symptom threshold). Using a less stringent control group, Chen and colleagues (2021)^14^ pooled results across 17 studies in their meta-analytic study (totaling 381 individuals with lifetime STBs compared to 642 healthy controls) and found hyperactivation in the bilateral superior temporal gyrus, left middle temporal gyrus, and right inferior parietal lobe and hypoactivation in the left insula and right cerebellum among individuals with lifetime suicide attempt(s) when compared to healthy controls.

Prevailing psychosocial theories of suicide conceptualize distinct etiologic factors leading to the development of suicidal thoughts versus behaviors^4, 6, 15^. Such theories posit that dispositional contributors, including biological factors like brain activation, may uniquely differentiate those at risk for suicidal behaviors versus suicidal thoughts alone^5^. Neuroimaging studies of suicide have lagged behind psychosocial research, as only a few studies to date have compared individuals with suicide attempt histories to those with other STBs^12^. In a small sub-analysis, 18 individuals with current suicidal ideation and lifetime suicide attempt(s) had marginally higher connectivity between the dorsal posterior cingulate cortex and the left inferior frontal gyrus compared to 16 individuals with current suicidal ideation without lifetime suicide attempt(s)^16^. Furthermore, lifetime suicide attempt(s), compared with suicidal ideation alone is associated with altered frontal brain function during tasks in several small imaging studies^17-19^.

Examination of altered functional networks may aid our understanding of suicide risk. Neuroscientists have increasingly recognized the brain is organized into intrinsic functional networks, or networks of regions that are commonly correlated or anticorrelated with each other at a given time^20-24^. As recommended by Uddin and colleagues (2019)^25^, an anatomical taxonomy should be used to refer to these networks rather than a functional taxonomy to enable greater reproducibility and consistency. For example, utilizing traditional cognitive nomenclature like “attention network” diminishes the role of these networks in other tasks and reduces reproducibility between research groups. Thus, an anatomical taxonomy will be emphasized in this paper (e.g., “dorsal frontoparietal network” rather than “attention network”).

The Triple Network Theory^26^ posits that three networks support the majority of cognitive and emotional processes and are central to clinically-concerning psychological dysfunction. First, the medial frontoparietal network (M-FPN; functionally referred to as the “default mode network”) consists of multiple smaller networks and includes parts of the mPFC, PCC, and hippocampus. This network is involved in self-referential processing, autobiographical memory retrieval, and future-oriented thinking^21, 27^.

Second, the lateral frontoparietal network (L-FPN; functionally referred to as the “cognitive/executive control network”) consists of lateral prefrontal regions along the middle frontal gyrus including the rostral and dorsolateral prefrontal cortex and the anterior inferior parietal lobule. This network is involved in goal-directed responses, emotion regulation, and some attentional processes^25, 28, 29^. Third, the mid-cingulo-insular network (M-CIN; functionally referred to as the “salience network”) consists of the dACC, bilateral anterior insula, and anterior midcingulate cortex. This network is involved in the detection of behaviorally relevant environmental stimuli and coordinating responses^25, 29^.

Two studies to date have directly tested the triple network theory in relation to suicide risk. Ordaz and colleagues (2018)^30^ examined the relationship between within-network M-FPN, L-FPN, and M-CIN functional connectivity among different brain regions within the same network and lifetime suicidal ideation severity (with no analyses pertaining to lifetime suicide attempts) in a sample of 40 adolescents diagnosed with major depressive disorder (MDD). Increased within-network functional connectivity in each of the three networks was independently associated with greater lifetime severity of suicidal ideation (i.e., longer duration, less controllability, higher frequency). Malhi and colleagues (2019)^31^ compared network connectivity of 25 individuals with mood disorders and lifetime suicide attempt(s) against 54 individuals with mood disorders and no lifetime suicide attempt(s) in each of the three networks and the basal ganglia network. Findings also revealed that increased posterior M-FPN activity was associated with past-month STBs, linking recent suicidality to default mode activity and potentially self-referential thinking^31^. Unfortunately, the lack of a suicidal ideation control group limits interpretations as M-FPN connectivity differences could be related to ideation, behavior, or both. While both studies represent important contributions, neither accounted for important potentially confounding variables or had sufficient sample size to directly examine between-network connectivity.

Only around 7% of imaging studies of suicide have compared individuals with lifetime suicide attempt(s) to controls with suicidal ideation alone^9^, which limits possible conclusions regarding the transition from suicidal ideation to behavior. Of those studies utilizing suicidal control groups, studies have largely been underpowered and lacked covariates. To address gaps in the scientific literature, we examined differences in resting-state functional brain network connectivity using a large subsample from the UK Biobank. Specifically, we compared individuals with lifetime suicide attempt(s) versus those with lifetime suicidal and/or non-suicidal self-injurious ideation (hereafter referred to as “self-directed violence thoughts,” SDVT per CDC nomenclature recommendations^32^) alone. The study aims were to compare resting-state connectivity both within (aim 1), and between (aim 2) M-FPN, L-FPN, and M-CIN network regions across the two study groups. In line with the prior research^26, 30, 31^, we hypothesized that individuals with lifetime suicide attempt(s) in comparison to those with lifetime SDVT alone would demonstrate (1) greater connectivity within the M-FPN and M-CIN networks but lower within-network connectivity among L-FPN regions, and (2) lower between-network connectivity among the M-FPN, L-FPN, and M-CIN network regions. Between-group differences in additional networks throughout the brain were additionally explored.

## Methods and Materials

### Participants

The UK Biobank is a population-based biomedical study of roughly 500,000 individuals from Great Britain (England, Scotland, and Wales) between the ages of 40 and 69^33^. Individuals enrolled in the UK Biobank answered demographic and medical questions and several weeks later, completed an online mental health follow-up questionnaire packet. Those who answered “yes” to the question “Have you deliberately harmed yourself, whether or not you meant to end your life?” (UK Biobank field 20480) were prompted to answer further detailed questions regarding “harm behaviours.” Among those questions were “Have you harmed yourself with the intention to end your life?” (suicide attempt history; UK Biobank field 20483) and “Have you contemplated harming yourself (for example, by cutting, biting, hitting yourself, or taking an overdose)?” (suicide-related thought history; UK Biobank field 20485). For the purposes of this study, this latter endorsement has been defined as “self-directed violence thoughts” (SDVT) in line with the U.S. Centers for Disease Control and Prevention (CDC) self-directed violence classification system^32^. A subsample of participants underwent magnetic resonance imaging (MRI) neuroimaging procedures, including structural and functional imaging^33^. Participants were excluded from MRI imaging if they reported neurological conditions/incidents^33^.

This study utilized data from a subsample of 4,013 individuals who either had a lifetime SDVT alone (*n* = 3,447) or in combination with suicide attempt(s) (*n* = 566) with valid functional neuroimaging data. A flowchart leading to the final study sample is depicted in Figure 1. Of those included in the final sample, 14.1% reported lifetime suicide attempt(s) and 85.9% reported lifetime SDVT alone.

**Figure 1.**
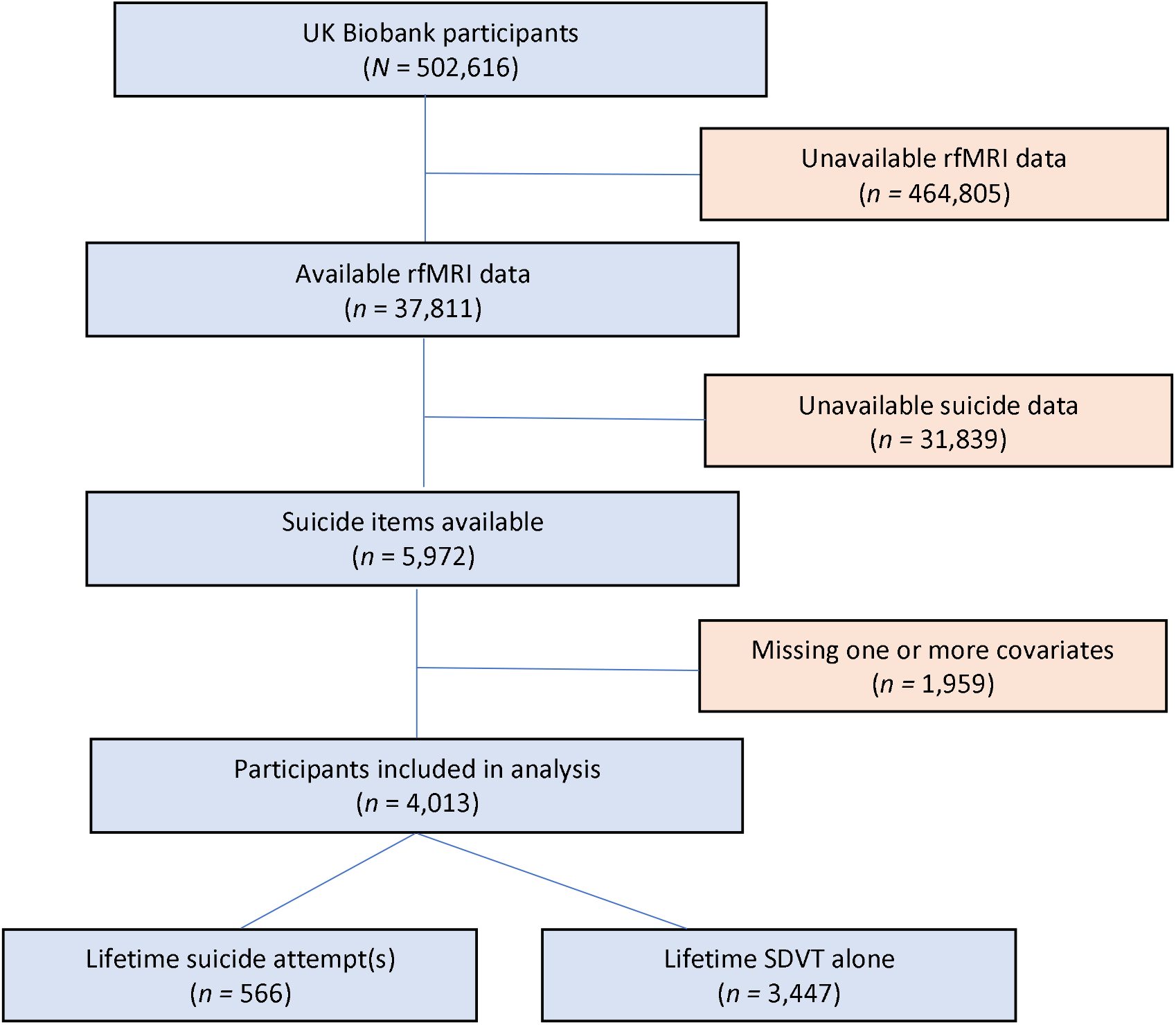
Flowchart of case selection.

### MRI Acquisition and Processing

MRI data were acquired in a Siemens Skya 3T scanner using a standard Siemens 32-channel head coil^33^. Briefly, 3D T1-weighted MPRAGE were acquired at 1×1×1 mm (208×256×256 field of view [FOV] matrix) and 2×2×2 (104×104×72 FOV matrix), respectively. Preprocessing was done using FSL tools by the UK Biobank team (https://fsl.fmrib.ox.ac.uk/fsl/fslwiki). Initial preprocessing included “defacing” for participant anonymity via linear transformation to mask out facial structures.

Preprocessing of T1 data included skull stripping, bias field correction, warping to MNI space using FNIRT^34^, and tissue-type segmentation using FAST^35^ to differentiate cerebrospinal fluid, gray, and white matter volumes and generate 139 image-derived phenotypes (IDPs). For further information, detailed UK Biobank data acquisition and preprocessing protocol (https://www.fmrib.ox.ac.uk/ukbiobank/protocol/V4_23092014.pdf) and associated documentation (http://biobank.ctsu.ox.ac.uk/crystal/docs/brain_mri.pdf) are freely available online.

For resting-state functional MRI (rsfMRI) procedures, UK Biobank participants were instructed to “keep their eyes fixated on a crosshair, relax, and think of nothing in particular.” Resting-state fMRI data were acquired using a resolution of 2.4×2.4×2.4 (88×88×64 FOV matrix) with TR = 0.735 s, TE = 39 ms, and GE-EPI with x8 multislice acceleration, no iPAT, flip angle 52? over 6 minutes (490 timepoints)^33^. Data preprocessing, group-independent components analysis (ICA) parcellation, and connectivity estimation were carried out by UK Biobank with FSL packages. These included motion correction with MCFLIRT^36^, grand-mean intensity normalization with a single multiplicative factor, high pass temporal filtering with a Gaussian-weighted least squares straight line fitting (sigma as 50.0 s), EPI unwarping using field map scanned before collection, gradient distortion correction (GDC) unwarping, and removal of structural artefacts using an ICA-based X-noiseifier^37^. Gross preprocessing failures were visually inspected by UK Biobank and removed^33^. Group-ICA parcellated preprocessed EPI images were fed into the MELODIC tool of FSL to generate a 21×21 matrix of ICA components, used for analyses (https://www.fmrib.ox.ac.uk/ukbiobank/protocol/V4_23092014.pdf)^37,38^.

### Analyses

Time series data from the 21 components were used for connectivity analysis, using each component as a node. A 21×21 partial matrix of fully-normalized partial temporal correlations were derived for each participant, as they represent direct connections better than full temporal correlations and control for the strength of other connections^37, 38^. For each component, a larger number indicates stronger temporal connectivity while positive or negative values represent valence. Prior to analysis, the strength of each connection was multiplied by the sign of its group mean^39^. This allowed for investigation of the degree to which temporal connectivity differed by history of suicide attempt without combining positive and negative effects and losing information about the absolute magnitude^37^.

The association between history of attempted suicide and the strength of connections was tested using the *glm* function in R, controlling for age, sex, ethnicity, and body mass index (BMI) based on the scientific literature^15, 37, 40-43^. BMI was calculated as weight (kg)/height^2^ (m). To compare within-network connectivity, 14 general linear regressions were performed comparing the two groups for within-network nodes representing the three networks of interest (one regression per pair within a given network). To compare between-network connectivity, 31 general linear regressions were performed comparing the two groups for between-network nodes representing the three networks of interest. To compare groups on additional brain networks,165 general linear regressions were performed comparing the two groups on the remaining 11 nodes (165 comparisons). False discovery rate (FDR) correction was applied over each set of tests (14 tests for within-network, 31 for between-network, and 165 for exploratory analyses) using the *p*.*adjust* function in R, setting *q* < 0.05 as the significance level^38^.

Sensitivity analyses were conducted evaluating Aims 1 and 2 without covariates. We have provided our scripts for conducting our analyses online (https://github.com/CNPsyLab/UKB-Suicide-Resting-State-Network-Analyses) to facilitate replication and extension of these findings with additional participants and novel analyses.

## Results

### Demographics

Data from a total of 4,013 individuals were included in analysis of neuroimaging correlates. Individuals included were on average 52.75 years old (SD = 7.13, range = 40-70 years old) at the time of initial study visit. Women represented 65.7% of the sample (*n* = 2,637) and 97.8% of the sample were non-Hispanic White. Those with lifetime suicide attempt(s) were more likely to be female, χ^*2*^ (1, 4013) = 5.97, *p* = 0.0146, compared with those with lifetime SDVT alone. Groups did not differ significantly based on age or ethnicity. Demographic characteristics for the overall sample and by group are presented in Table 1.

**Table 1.** Demographic characteristics for the overall sample and by group.

| | Overall<br>(*n* = 4,013) | | Group 1: SA<br>(*n* = 566) | | Group 2: SDVT<br>(*n* = 3,447) | | $\chi^2$ | *p* |
| --- | --- | --- | --- | --- | --- | --- | --- | --- |
|  | Frequency | Percent | Frequency | Percent | Frequency | Percent |  |  |
| Age, Mean (SD), *t* statistic | 52.75 (range<br>= 40-70) | (7.13) | 52.90 | (6.97) | 52.90 | (7.16) | -0.02 | 0.9807 |
| Sex |  |  |  |  |  |  |  |  |
| Female | 2637 | 65.7% | 398 | 70.3% | 2239 | 65.0% | **5.97** | **0.0146** |
| Male | 1376 | 34.3% | 168 | 29.7% | 1208 | 35.0% |  |  |
| Race/Ethnicity† |  |  |  |  |  |  |  |  |
| Non-Hispanic White | 3925 | 97.8% | 556 | 98.2% | 3369 | 97.7% | 0.49 | 0.4833 |
| Non-White | 79 | 2.0% | 9 | 1.6% | 70 | 2.0% |  |  |
| Prefer not to answer | 7 | 0.2% | 0 | 0% | 7 | 0.2% |  |  |
| Do not know | 1 | 0% | 1 | 0.2% | 0 | 0% |  |  |
| BMI, Mean (SD), *t* statistic | 26.62 | (4.62) | 27.03 | (4.98) | 26.55 | (4.56) | **-2.18** | **0.0293** |

Note. Data reported as *n* (%), unless otherwise specified. Group 1 (SA) = Lifetime Suicide Attempt(s); Group 2 (SDVT) = Lifetime Self-Directed Violence Thoughts (SDVT) Alone.
†One participant in Group 2 was missing data on race/ethnicity and was coded as “prefer not to answer” for subsequent analyses. Due to expected counts <5, race/ethnicity chi-square analysis was conducted using only Non-Hispanic White and Non-White categories without “Prefer not to answer” or “Do not know.”

#### Within-network Connectivity

As shown in Table 2, no models revealed statistically significant differences between groups after adjusting for multiple corrections (FDR *q* > 0.05). Subsequent sensitivity analyses conducted without covariates similarly did not reveal statistically significant differences between groups after adjusting for multiple corrections (FDR *q* > 0.05).

**Table 2.** Comparisons of two selected groups on within-network resting-state connectivity among the medial frontoparietal, lateral frontoparietal, and midcingulo-insular networks.

| Network | Network 1<br>(Node) | Network 2<br>(Node) | *t*-value | SE | $\beta$ | *p*-value | FDR-adjusted<br>*q*-value |
| --- | --- | --- | --- | --- | --- | --- | --- |
| *Medial*<br>*Frontoparietal* | M-FPN (1) | M-FPN (7) | -2.04 | 0.04 | -0.03 | 0.0419* | 0.2217 |
|  | M-FPN (1) | M-FPN (9) | 1.00 | 0.04 | 0.02 | 0.3180 | 0.7420 |
|  | M-FPN (1) | M-FPN (14) | -0.61 | 0.04 | -0.01 | 0.5404 | 0.8406 |
|  | M-FPN (1) | M-FPN (20) | 1.70 | 0.04 | 0.03 | 0.0894 | 0.3132 |
|  | M-FPN (7) | M-FPN (9) | -0.808 | 0.04 | -0.01 | 0.4189 | 0.8378 |
|  | M-FPN (7) | M-FPN (14) | 0.33 | 0.03 | 0.01 | 0.7430 | 0.9378 |
|  | M-FPN (7) | M-FPN (20) | 0.02 | 0.04 | <0.01 | 0.9816 | 0.9828 |
|  | M-FPN (9) | M-FPN (14) | 0.16 | 0.03 | <0.01 | 0.8697 | 0.9828 |
|  | M-FPN (9) | M-FPN (20) | -0.70 | 0.04 | -0.01 | 0.4825 | 0.8406 |
|  | M-FPN (14) | M-FPN (20) | 2.11 | 0.03 | 0.03 | 0.0354* | 0.2217 |
| *Lateral*<br>*Frontoparietal* | L-FPN (5) | L-FPN (16) | 0.02 | 0.04 | <0.01 | 0.9828 | 0.9828 |
| *Midcingulo-*<br>*Insular* | M-CIN (13) | M-CIN (18) | -1.37 | 0.02 | -0.02 | 0.1702 | 0.4766 |
|  | M-CIN (13) | M-CIN (21) | -1.98 | 0.04 | -0.03 | 0.0475* | 0.2217 |
|  | M-CIN (18) | M-CIN (21) | 0.04 | <0.01 | <0.01 | 0.9724 | 0.9828 |

Note. Group 1 (SA) = Lifetime Suicide Attempt(s); Group 2 (SDVT) = Lifetime Self-Directed Violence Thoughts (SDVT) Alone; BMI = body mass index; SE = standard error; FDR = false discovery rate; M-FPN = medial frontoparietal network; L-FPN = lateral frontoparietal network; M-CIN = midcingulo-insular network. All models adjusted for age, sex, ethnicity, and BMI

#### M-FPN Within-network Connectivity

Prior to FDR correction for multiple comparisons, two of the ten models showed altered connectivity within the M-FPN. Participants with history of suicide attempt(s) had lower connectivity between node 1 and node 7 in comparison to those with a history of SDVT alone, *t*(1, 4013) = -2.04, *p* = 0.0419, FDR *q* = 0.2217. These nodes included connectivity between areas of the ventromedial prefrontal cortex (node 1) with areas of the retrosplenial and medial temporal cortices (node 7). Participants with lifetime suicide attempt(s) had greater connectivity between node 14 and node 20 in comparison to those lifetime SDVT alone, *t*(1, 4013) = 2.11, *p* = 0.0354, FDR *q* = 0.2217. These nodes include connectivity between areas of the anterior cingulate and orbitofrontal cortices (node 14) with areas of the posterior precuneus and posterior cingulate cortex (node 20). Figure S1 depicts significant between-network group differences prior to FDR correction.

#### L-FPN Within-network Connectivity

One model examined group differences on connectivity between nodes associated with the L-FPN. As shown in Table 2, this model did not reveal statistically significant differences between groups.

#### M-CIN Within-network Connectivity

Prior to FDR correction for multiple comparisons, one of the three models showed differences in connectivity within the M-CIN. Individuals with lifetime suicide attempt(s) had lower connectivity between node 13 and node 18 in comparison to those with lifetime SDVT alone, *t*(1, 4013) = -1.98, *p* = 0.0475, FDR *q* = 0.2217. As depicted in Figure 2c, these nodes include connectivity between areas of the left cingulo-opercular cortex (node 13) with areas of the putamen, striatum, and basal ganglia (node 18).

### Between-network Connectivity

As shown in Table 3, no models revealed statistically significant differences between groups after adjusting for multiple corrections (FDR *q* > 0.05). Subsequent sensitivity analyses conducted without covariates similarly did not reveal statistically significant differences between groups after adjusting for multiple corrections (FDR *q* > 0.05).

**Table 3.**
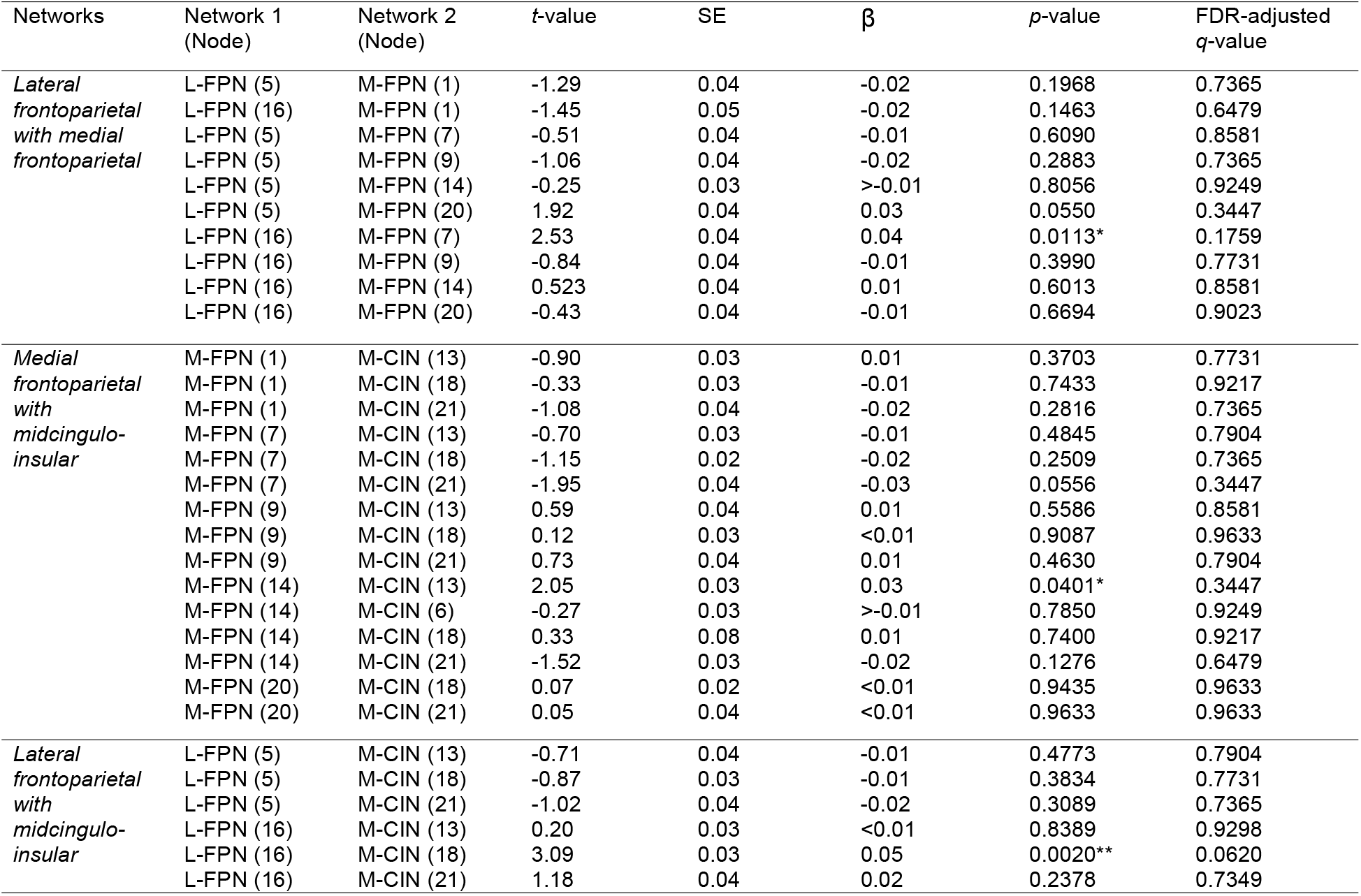
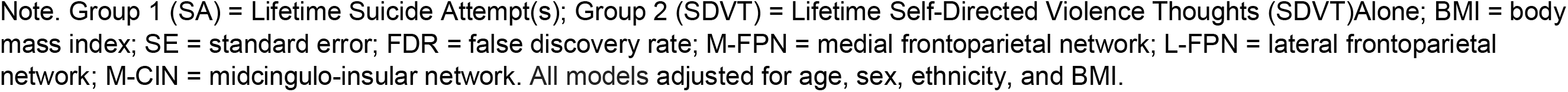
Comparisons of two selected groups on between-network resting-state connectivity among the medial frontoparietal, lateral frontoparietal, and midcingulo-insular networks.

| Networks | Network 1 (Node) | Network 2 (Node) | t-value | SE | $\beta$ | p-value | FDR-adjusted q-value |
| --- | --- | --- | --- | --- | --- | --- | --- |
| *Lateral frontoparietal with medial frontoparietal* | L-FPN (5) | M-FPN (1) | -1.29 | 0.04 | -0.02 | 0.1968 | 0.7365 |
|  | L-FPN (16) | M-FPN (1) | -1.45 | 0.05 | -0.02 | 0.1463 | 0.6479 |
|  | L-FPN (5) | M-FPN (7) | -0.51 | 0.04 | -0.01 | 0.6090 | 0.8581 |
|  | L-FPN (5) | M-FPN (9) | -1.06 | 0.04 | -0.02 | 0.2883 | 0.7365 |
|  | L-FPN (5) | M-FPN (14) | -0.25 | 0.03 | >-0.01 | 0.8056 | 0.9249 |
|  | L-FPN (5) | M-FPN (20) | 1.92 | 0.04 | 0.03 | 0.0550 | 0.3447 |
|  | L-FPN (16) | M-FPN (7) | 2.53 | 0.04 | 0.04 | 0.0113* | 0.1759 |
|  | L-FPN (16) | M-FPN (9) | -0.84 | 0.04 | -0.01 | 0.3990 | 0.7731 |
|  | L-FPN (16) | M-FPN (14) | 0.523 | 0.04 | 0.01 | 0.6013 | 0.8581 |
|  | L-FPN (16) | M-FPN (20) | -0.43 | 0.04 | -0.01 | 0.6694 | 0.9023 |
| *Medial frontoparietal with midcingulo-insular* | M-FPN (1) | M-CIN (13) | -0.90 | 0.03 | 0.01 | 0.3703 | 0.7731 |
|  | M-FPN (1) | M-CIN (18) | -0.33 | 0.03 | -0.01 | 0.7433 | 0.9217 |
|  | M-FPN (1) | M-CIN (21) | -1.08 | 0.04 | -0.02 | 0.2816 | 0.7365 |
|  | M-FPN (7) | M-CIN (13) | -0.70 | 0.03 | -0.01 | 0.4845 | 0.7904 |
|  | M-FPN (7) | M-CIN (18) | -1.15 | 0.02 | -0.02 | 0.2509 | 0.7365 |
|  | M-FPN (7) | M-CIN (21) | -1.95 | 0.04 | -0.03 | 0.0556 | 0.3447 |
|  | M-FPN (9) | M-CIN (13) | 0.59 | 0.04 | 0.01 | 0.5586 | 0.8581 |
|  | M-FPN (9) | M-CIN (18) | 0.12 | 0.03 | <0.01 | 0.9087 | 0.9633 |
|  | M-FPN (9) | M-CIN (21) | 0.73 | 0.04 | 0.01 | 0.4630 | 0.7904 |
|  | M-FPN (14) | M-CIN (13) | 2.05 | 0.03 | 0.03 | 0.0401* | 0.3447 |
|  | M-FPN (14) | M-CIN (6) | -0.27 | 0.03 | >-0.01 | 0.7850 | 0.9249 |
|  | M-FPN (14) | M-CIN (18) | 0.33 | 0.08 | 0.01 | 0.7400 | 0.9217 |
|  | M-FPN (14) | M-CIN (21) | -1.52 | 0.03 | -0.02 | 0.1276 | 0.6479 |
|  | M-FPN (20) | M-CIN (18) | 0.07 | 0.02 | <0.01 | 0.9435 | 0.9633 |
|  | M-FPN (20) | M-CIN (21) | 0.05 | 0.04 | <0.01 | 0.9633 | 0.9633 |
| *Lateral frontoparietal with midcingulo-insular* | L-FPN (5) | M-CIN (13) | -0.71 | 0.04 | -0.01 | 0.4773 | 0.7904 |
|  | L-FPN (5) | M-CIN (18) | -0.87 | 0.03 | -0.01 | 0.3834 | 0.7731 |
|  | L-FPN (5) | M-CIN (21) | -1.02 | 0.04 | -0.02 | 0.3089 | 0.7365 |
|  | L-FPN (16) | M-CIN (13) | 0.20 | 0.03 | <0.01 | 0.8389 | 0.9298 |
|  | L-FPN (16) | M-CIN (18) | 3.09 | 0.03 | 0.05 | 0.0020** | 0.0620 |
|  | L-FPN (16) | M-CIN (21) | 1.18 | 0.04 | 0.02 | 0.2378 | 0.7349 |

Note. Group 1 (SA) = Lifetime Suicide Attempt(s); Group 2 (SDVT) = Lifetime Self-Directed Violence Thoughts (SDVT) Alone; BMI = body mass index; SE = standard error; FDR = false discovery rate; M-FPN = medial frontoparietal network; L-FPN = lateral frontoparietal network; M-CIN = midcingulo-insular network. All models adjusted for age, sex, ethnicity, and BMI.

#### L-FPN with M-FPN

Prior to FDR corrections for multiple comparisons, one of ten models showed altered connectivity between nodes associated with the L-FPN with nodes associated with the M-FPN. Individuals with lifetime suicide attempt(s) had increased connectivity between node 16 (L-FPN) and node 7 (M-FPN) in comparison to those with lifetime SDVT alone, *t*(1, 4013) = 2.53, FDR *q* > 0.05. These nodes include connectivity between areas of the anterior and dorsolateral prefrontal cortex (node 16) with areas of the retrosplenial and medial temporal cortices (node 7).

#### M-FPN with M-CIN

Prior to FDR corrections for multiple comparisons, one of fifteen models showed altered connectivity between nodes associated with the M-FPN with nodes associated with the M-CIN. Individuals with lifetime suicide attempt(s) had greater connectivity between node 13 and node 14 in comparison to those with lifetime SDVT alone, *t*(1, 4013) = 2.05, *p* = 0.0401, FDR *q* = 0.3447. These nodes include connectivity between areas of the left cingulo-opercular cortex (node 13) with areas of the anterior cingulate and orbitofrontal cortices (node 14).

#### L-FPN with M-CIN

Prior to FDR corrections for multiple comparisons, one of six models examined group differences on connectivity between nodes associated with the L-FPN with nodes associated with the M-CIN. Participants with lifetime suicide attempt(s) had greater connectivity between node 16 (L-FPN) and node 18 (M-CIN) in comparison to those with lifetime SDVT alone, *t*(1, 4013) = 3.09, *p* = 0.0020, FDR *q* = 0.0620. These nodes include connectivity between areas of the anterior and dorsolateral prefrontal cortex (node 16) with areas of the putamen, striatum, basal ganglia, and thalamus (node 18). Figure S2 depicts significant between-network group differences prior to FDR correction.

### Whole-brain Connectivity

No models revealed statistically significant differences between groups after adjusting for multiple corrections (FDR *q* > 0.05).

## Discussion

In the largest functional imaging study of suicide behavior to date, we compared resting-state connectivity both within and between M-FPN, L-FPN, and M-CIN network regions among individuals with lifetime suicide attempt(s) versus those with lifetime SDVT alone. No significant between-group differences were found after correcting for multiple comparisons. Specifically, we found no significant group differences in within-or between-network connectivity among nodes of the M-FPN, L-FPN, or M-CIN. Further, there were no significant group differences on exploratory whole-brain connectivity analyses.

Despite its sample size powered to detect small effects, this study did not find significant within-network, between-network, or whole-brain connectivity differences after correcting for multiple comparisons. These results contrast with several smaller studies that previously found resting-state differences between those with lifetime suicide attempt(s) and psychiatric controls^44-46^ but are consistent with pooled findings in meta-analyses^9, 14^. Lack of findings supporting hypotheses are consistent with growing trends in brain science research showing reduced effects upon replication in larger samples^47^. Functional neuroimaging studies have, in particular, been underpowered^48, 49^, with an inverse relationship between sample size and number of significant findings. The average sample size of neuroimaging studies of suicide risk is around 48^9^, which indicates that many previously found differences may be inflated or spurious.

Null findings in the present study may suggest a more complex relationship between dispositional factors of suicide risk, like brain circuitry, and the transition from suicide-related thoughts to behaviors. Rather, the ideation-to-action theoretical framework would necessitate that an examination of the complex interaction among biopsychosocial factors – more specifically, dispositional (biological, genetic), acquired (learning), and practical aspects (knowledge of and access to lethal means)^5^ – is needed to fully understand how individuals with lifetime self-directed violence thoughts alone may differ from those who eventually progress to suicide attempts.

With regards to within-network connectivity, there was limited evidence to support differences between individuals with lifetime suicide attempt(s) versus those with lifetime SDVT alone after correcting for multiple comparisons. Malhi and colleagues (2019) found within-network connectivity differences in the M-FPN when comparing those with attempted suicide to healthy controls, but did not find differences when those with attempted suicide were compared to those diagnosed with a mood disorder^31^.

Similarly, a recent study comparing 35 depressed adolescents with lifetime suicide attempt(s) to 18 adolescents with mood disorder without lifetime suicide attempt(s) did not find within-network differences when looking at regions within the M-FPN and M-CIN^50^. This suggests that within-network differences may be too subtle to detect when using psychiatric or SDVT controls.

With regards to between-network connectivity, this study did not find significant differences between-group differences after correcting for multiple comparisons. This is consistent with Mahli and colleagues (2019) who did not find between-network connectivity when comparing those with attempted suicide to healthy controls or individuals diagnosed with a mood disorder. In their meta-analysis, Huang and colleagues (2020)^9^ found that, compared to all controls (healthy and psychiatric), those with prior suicidal ideation and behaviors collectively showed hyperactivation of the right posterior cingulate cortex and superior frontal gyrus during pooled affective tasks, suggesting potential alterations between the M-FPN and L-FPN during affective processes. In similar meta-analysis of functional imaging studies, Chen and colleagues (2021)^14^ found hyperactivation of the bilateral superior temporal gyrus in pooled studies among those with suicide attempt compared to all controls. Larger studies are needed to more fully investigate potential between-network differences during cognitive, affective, and social tasks.

Finally, this study did not find other significant whole-brain connectivity between-group differences. Results are consistent with a recent meta-analyses of event-related potential studies, which found small or no effects in larger studies comparing those with suicide attempt to suicidal ideation alone^51^. Null findings across methodologies using stringent control groups may reflect the need for more nuanced investigation and interpretation of the interplay between biopsychosocial factors to differentiate those with lifetime history of suicide attempt(s) from SDVT alone^5^. Differentiating factors uniquely associated with suicide attempt has been a challenge within the field of suicidology^52^.

Covariate use in neuroimaging studies is highly diverse, with some studies reporting use of zero covariates and others using as many as 14 covariates in analyses^53^. Smith, Nichols ^40^ highlight how large datasets, in particular, are especially susceptible to artifactual associations due to confounding effects in neuroimaging research. Alfaro-Almagro and colleagues (2020) caution careful consideration of covariates, particularly in relation to IDP imaging data in the UK Biobank. Both research groups suggest motion correction and noise removal to reduce imaging-related artifacts in analysis, both of which were conducted prior to these analyses. Further, near duplication of findings after sensitivity analyses in this study conducted without covariates increases confidence that lack of robust findings was not due to covariate selection.

There are several noteworthy limitations to this study. Without a comprehensive suicide risk assessment to reveal lifetime STBs using robust instruments such as the Columbia Suicide Severity Rating Scale, it is difficult to discern the temporal relationship between suicide attempt history and functional connectivity differences. Individuals examined in this study were 53 years old, on average, thus suicide attempt(s) and SDVT could be distal events. Functional markers may be better than structural anatomical markers at clinically differentiating those who are acutely suicidal in the future, as suicide thoughts are typically time-limited^54, 55^. Future studies can incorporate time since suicide attempt(s) and SDVT to account for potential differences in time.

Relatedly, without a comprehensive lifetime suicide history and risk assessment, there are several confounding variables. Using a SDVT control group, we were unable to parse those with lifetime suicidal versus non-suicidal self-directed violence ideation. While a SDVT control group is a strength compared to previous studies, there may be important distinctions between those who have thoughts of suicidal (e.g., thinking about using a firearm to kill oneself) versus non-suicidal (e.g., thinking about non-lethal cutting or scratching) self-directed violence^56^. Further, we were unable to differentiate among individuals with single versus multiple suicide attempts. Those with multiple lifetime attempts have distinct features, including impulsivity and borderline personality disorder traits and associated symptoms, that may reflect differences in functional neuroimaging markers^57^. We are similarly limited in survivorship bias and cannot extend findings to those who have died by suicide. Though our study used a stringent control group, there may be variability within functional connectivity within suicidal groups. Future studies can advance our understanding of potential differences among individuals with lifetime suicidal versus non-suicidal self-directed violent ideation, as well as single versus multiple suicide attempts.

Despite these limitations, strengths of this study include its sample size, use of a SDVT control group, and corrections for multiple comparisons to avoid spurious findings. This study utilized empirically-supported anatomical markers for network nodes, IDP markers for network connectivity which have been validated in previous studies^37, 38^, and comparison of groups were based on both theory-(within- and between-network comparisons) and data-driven (whole-brain comparisons) outcomes. This study on within- and between-network connectivity in those with lifetime suicide attempt(s) adds to the growing literature of biological correlates of suicide risk.

## Conclusions

In the largest neuroimaging study examining suicide attempts, individuals with lifetime suicide attempt(s), when compared to those with lifetime self-directed violent thoughts alone, did not demonstrate within- or between-network connectivity differences in the M-FPN, L-FPN, M-CIN or other subnetworks after controlling for multiple comparisons. Findings highlight the need for well-powered neuroimaging studies of suicide behavior using stringent control groups. Dispositional risk factors, like those measured by functional neuroimaging, may be less straightforward and rather may interact with psychosocial risk factors (e.g., access to means) in differentiating those at risk for SDVT from suicide attempt(s). Overall, this provides support for further study into the complex relationship between brain function and suicidality.

