## Supplemental Figures for "Resting-State Network Analysis of Suicide Attempt History in the UK Biobank"

Figure S1. Lifetime suicide attempt(s) was nominally associated with altered within-network connectivity in M-FPN when compared with lifetime SDVT alone prior to FDR correction for multiple comparisons. Those with a history of suicide attempt had lower connectivity between M-FPN nodes 1 and 7 (blue line) and greater connectivity between M-FPN nodes 14 and 20 (red line). Lifetime suicide attempt(s) was also associated with lower connectivity between M-CIN nodes 13 and 21 (blue line) prior to FDR correction for multiple comparisons. Models were adjusted for age, sex, ethnicity, and BMI.

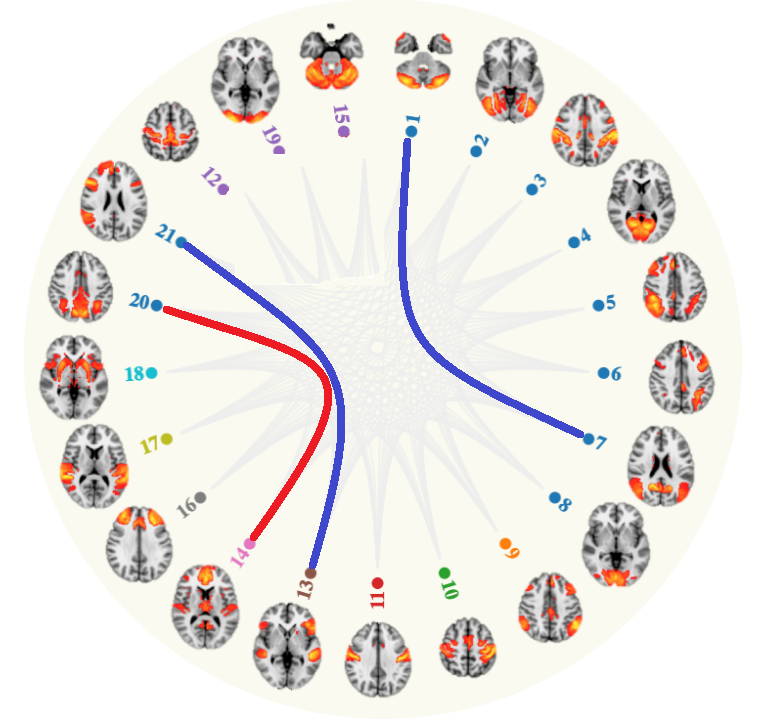

Figure S2. Lifetime suicide attempt(s) was nominally associated with stronger connectivity (red lines) between the lateral frontoparietal network (node 16) and medial frontoparietal network (node 7) and midcingulo-insular network (node 18) when compared with lifetime SDVT alone prior to FDR correction for multiple comparisons. Lifetime suicide attempt(s) was also associated with greater connectivity between the medial frontoparietal network (node 14) and the midcingulo-insular network (node 13). Models were adjusted for age, sex, ethnicity, and BMI.

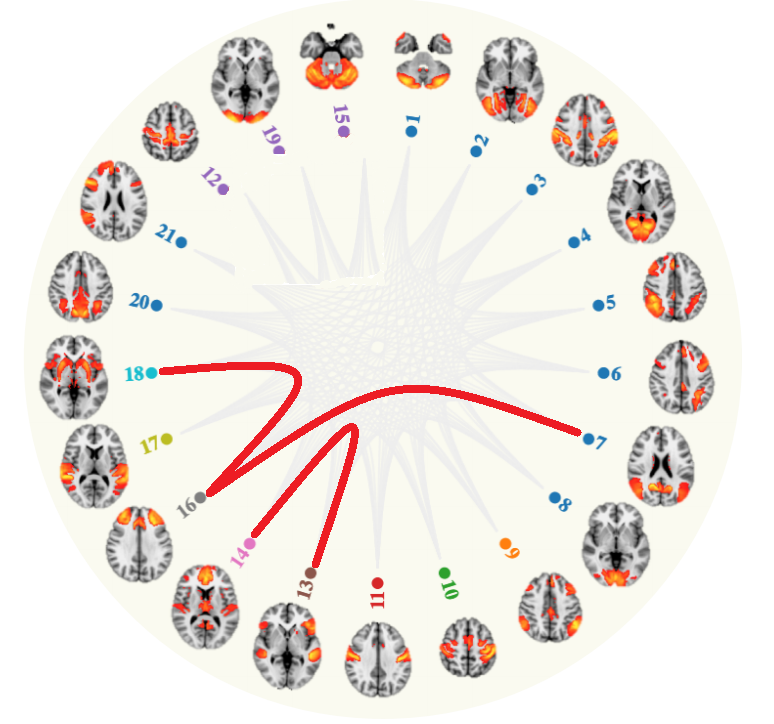
